# Harnessing changes in open chromatin determined by ATAC-seq to generate insulin-responsive reporter constructs

**DOI:** 10.1101/2021.05.06.443010

**Authors:** Collin B. Merrill, Austin B. Montgomery, Miguel A. Pabon, Aylin R. Rodan, Adrian Rothenfluh

## Abstract

**Background:** Gene regulation is critical for proper cellular function. Next-generation sequencing technology has revealed the presence of regulatory networks that regulate gene expression and essential cellular functions. Studies investigating the epigenome have begun to uncover the complex mechanisms regulating transcription. Assay for transposase-accessible chromatin by sequencing (ATAC-seq) is quickly becoming the assay of choice for many epigenomic investigations. However, whether intervention-mediated changes in accessible chromatin determined by ATAC-seq can be harnessed to generate intervention-inducible reporter constructs has not been systematically assayed.

**Results:** We used the insulin signaling pathway as a model to investigate chromatin regions and gene expression changes using ATAC- and RNA-seq in insulin-treated *Drosophila* S2 cells. We found correlations between ATAC- and RNA-seq data, especially when stratifying differentially-accessible chromatin regions by annotated feature type. In particular, our data demonstrated a strong correlation between chromatin regions annotated to distal promoters (1-2 kb from the transcription start site) and downstream gene expression. We cloned candidate distal promoter regions upstream of luciferase and demonstrate insulin-inducibility of several of these reporters.

**Conclusions:** Insulin-induced chromatin accessibility determined by ATAC-seq reveals enhancer regions that drive insulin-inducible reporter gene expression.

## Background

Gene regulation is essential to the development and maintenance of life. Gene regulatory networks describe the interplay between regulatory regions, such as promoters and enhancers, and expression of their target genes [1]. Deciphering how specific regulatory regions control gene transcription can provide insights into biological processes such as cell type differentiation [2, 3], responses to addictive substances [4], and other cell functions. The advent of new sequencing techniques has led to a greater understanding of how genes are differentially expressed. RNA-seq has provided a broader and more detailed picture of complex transcriptional states and responses [5, 6]. While genome-wide RNA-seq experiments can yield information on the many genes that are differentially transcribed in different conditions, these rich datasets reveal little about the regulatory mechanisms involved in directing these expression changes.

Epigenomic assays such as chromatin immunoprecipitation (ChIP-seq), DNAse-seq, and assay for transposase-accessible chromatin by sequencing (ATAC-seq) can interrogate chromatin accessibility and identify transcription factor binding sites [7–9]. The relationship between chromatin accessibility and transcription is complicated. Previous studies show little overlap between corresponding differences in chromatin and transcription [10–12], which highlights the complex interactions between the chromatin state and downstream gene expression. Furthermore, few studies have analyzed if changes in open chromatin induced by an intervention occur in transcriptional enhancers that can be coupled to heterologous minimal promoters to engineer intervention-inducible reporter constructs.

Here, we sought to characterize the relationship between ATAC-seq and RNA-seq data in more detail, with particular focus on whether intervention-induced changes in chromatin accessibility can accurately predict gene expression. We used the insulin signaling pathway as a model because the insulin receptor activates multiple downstream signaling pathways [13– 15] resulting in widespread changes to the chromatin state [16] and gene expression [17]. Our data from *Drosophila* S2 cells show that ATAC-seq and RNA-seq datasets are correlated, mainly driven by the ATAC-seq peaks/reads located in gene promoter regions. We also show that DNA regions with increased accessibility after insulin treatment can be harnessed to generate insulin-inducible reporter constructs.

## Results

### ATAC-seq and RNA-seq changes in insulin-exposed S2 cells

To investigate the concordance in changes in gene expression and chromatin accessibility, we exposed serum-starved *Drosophila* S2 cells to insulin or vehicle and harvested the cells 4 hours later for ATAC-seq and RNA-seq analysis. We determined genome-wide changes in open chromatin by ATAC-seq and identified 9726 high-confidence peaks (i.e., regions of accessible DNA mapped to the nuclear genome) in the insulin-exposed S2 cells, and 9560 in the vehicle-exposed S2 cells. Merging the control and experimental peak sets resulted in 10269 peaks. The largest variance in this dataset (6 samples; 2 treatments x 3 replicates) arose from insulin treatment, as shown by principal component analysis (PCA; Fig. 1A). In parallel, we identified 10287 transcripts in vehicle- and insulin-exposed S2 cells using RNA-seq. PCA indicated that the largest variance between the 6 samples resulted from insulin treatment (Fig. 1B). Because ATAC-seq provides a view of chromatin accessibility along all features of genes, we evaluated the feature distribution of both the treatment and the control ATAC-seq data (Fig. 1C). We observed the same genome features in the control and treated data, but the relative proportion of features was significantly different (χ^2^ = 19.6, df = 10, *p* = 0.03). This difference largely resulted from a change in the proportion of peaks annotated to distal (1-2 kb from the TSS) and proximal (≤1 kb) promoters, which increased from 8% to 9% and 58% to 60%, respectively. These results suggest that insulin signaling recruits additional regulatory features by changing chromatin accessibility.

**Figure 1.**
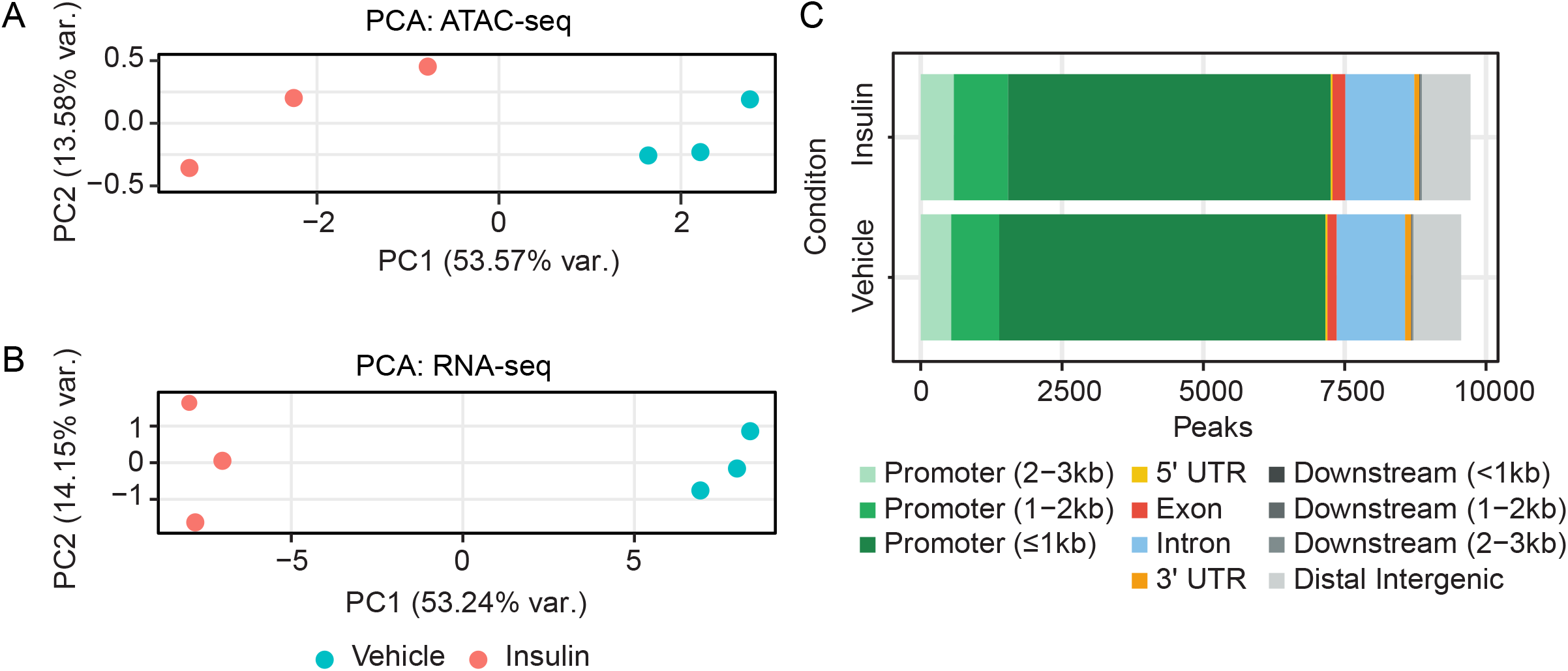
Insulin induces widespread alterations in chromatin accessibility and transcription. Serum-starved S2 cells were treated with vehicle or insulin for 4 h, and nucleic acids were isolated and analyzed. A) Principal component analysis of chromatin accessibility determined by ATAC-seq. B) Principal component analysis of transcript expression by RNA-seq. C) Proportions of each genomic feature type in all annotated chromatin peaks analyzed by ATAC-seq in S2 cells after treatment with insulin or vehicle.

### ATAC-seq and RNA-seq reads show weak correlation driven by ATAC-seq peaks in proximal promoters

We next asked whether RNA transcript levels were correlated with ATAC-seq reads and whether the feature annotation of those ATAC-seq peaks, i.e. where in a gene they were located, mattered for any levels of correlation. 8621 out of 10269 ATAC-seq peaks were mapped to a gene, and we plotted these ATAC-seq peak reads against the RNA-seq reads for each peak (thus duplicating many RNA-seq data points, since each gene has a median number of 2 (Quartile1-3: 2-4) ATAC-seq peaks mapped to it. Overall, RNA-seq and ATAC-seq peak reads showed a highly significant (*p* = 2.2e-16), but weak correlation (Pearson correlation coefficient *R* = 0.1; Fig 2A). When we stratified this analysis by the 11 ATAC-seq peak gene features, only the peak reads in the ≤1 kb promoter class correlated with RNA-seq reads (*R* = 0.2, *p* = 2.2e-16; Fig. 2B). This would suggest that more highly transcribed genes require a greater extent of DNA accessibility in their promoters, which might be expected for efficient transcriptional initiation.

**Figure 2.**
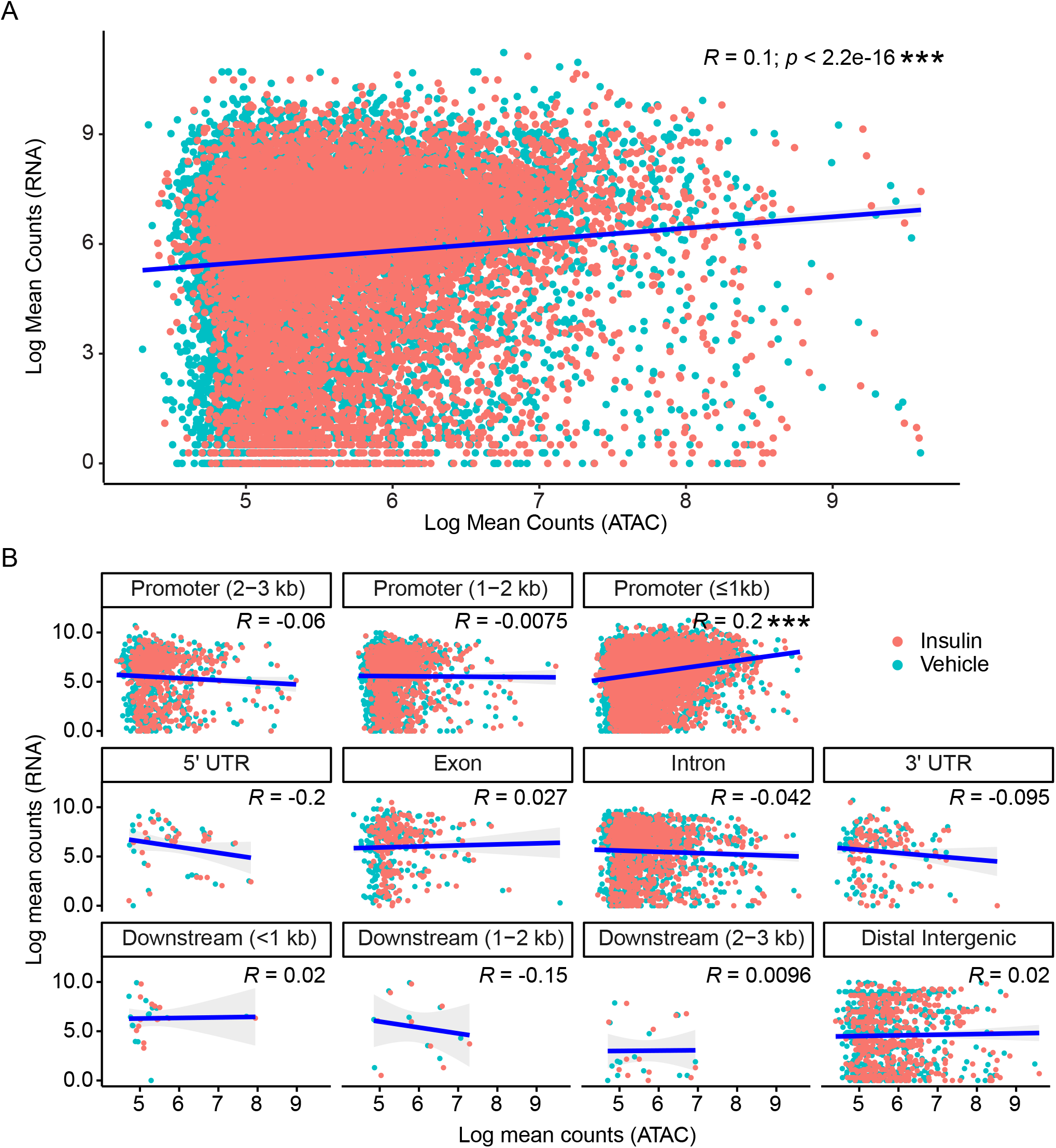
Chromatin peaks annotated to proximal promoters drive the correlation between normalized ATAC-seq and RNA counts. A) Transcripts identified by RNA-seq were overlapped with chromatin peaks annotated to the same genes. The normalized ATAC- and RNA-seq counts were log scaled and analyzed using Pearson correlation analysis. B) Overlapping ATAC- and RNA-seq counts from (A) were stratified by genomic feature. Pearson correlation analysis was used to identify feature-specific correlations between ATAC- and RNA-seq counts. Here, and in following figures, ns = not significant, \**p* < 0.05; \*\**p* < 0.01; \*\*\**p* < 0.001.

### Differential gene expression and DNA accessibility correlate for multiple ATAC-seq peak feature annotations

Next, we determined the insulin-induced changes in DNA accessibility and RNA expression. In the ATAC-seq peak set, 773 peaks were significantly differentially accessible (false discovery rate, FDR < 0.1) between the insulin-exposed and the control samples. 364 of the peaks were more accessible upon insulin exposure, while 409 peaks were less accessible after exposure to insulin (Fig. 3A). The feature distribution of those differential peaks was very similar to the feature distribution in the whole ATAC-seq peak dataset (χ^2^ = 6.13, df = 10, *p* = 0.80; Fig. 3B), though we did not detect distal downstream elements (1-2 and 2-3 kb downstream) in the differentially accessible peaks. We also examined the significant gene expression changes from the RNA-seq dataset. In this dataset, 3616 genes were differentially expressed (FDR < 0.05) between the insulin-exposed and control samples. 2056 genes were upregulated after insulin exposure, while 1560 were downregulated (Fig. 3C).

**Figure 3.**
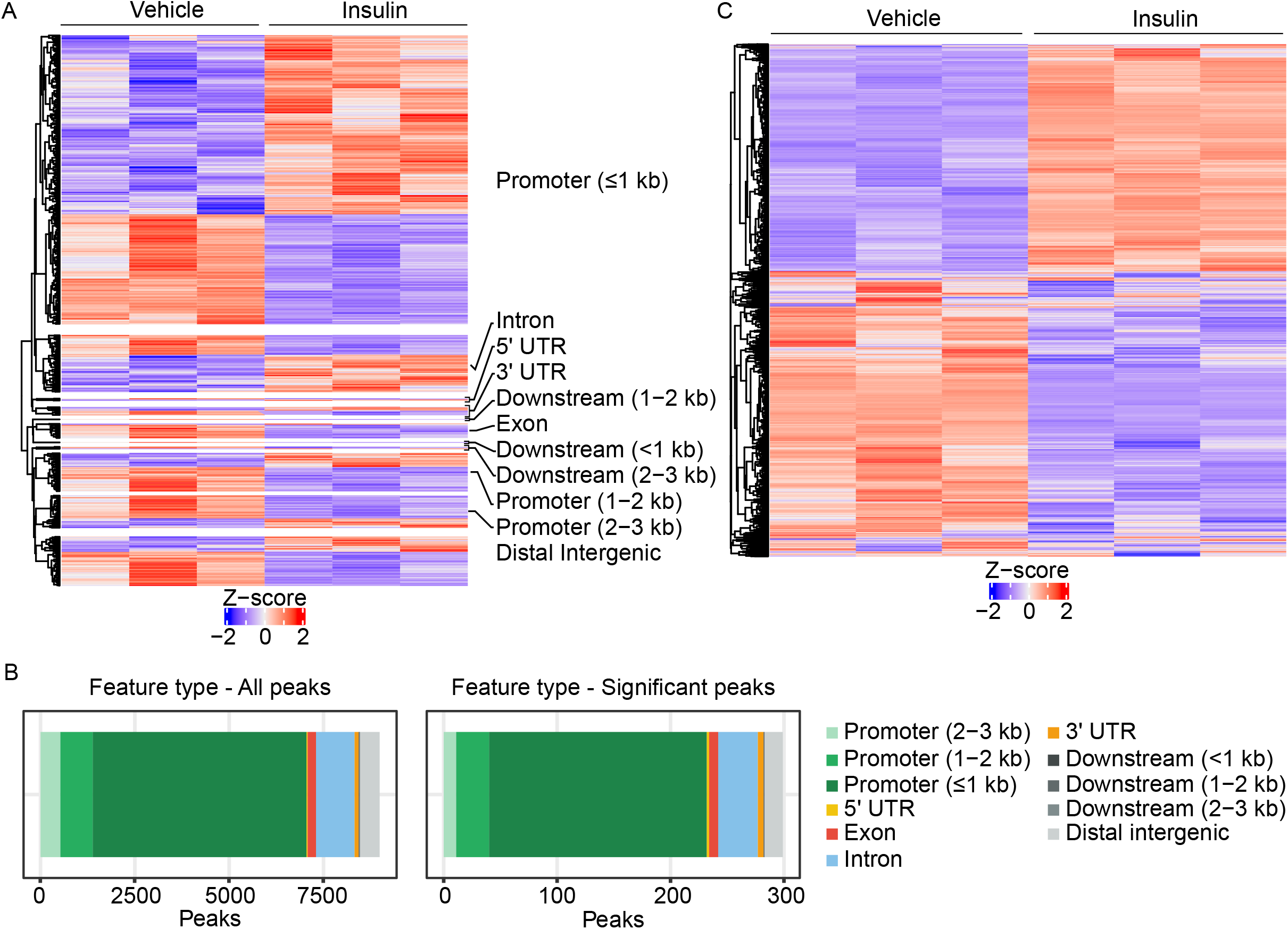
Insulin induces differential chromatin accessibility and gene expression in S2 cells. A) Heatmap of differential chromatin accessibility in significantly different chromatin peaks, stratified by feature type. Each row represents an individual chromatin peak. The values in each row were scaled to the row mean. Red indicates more-accessible chromatin regions and blue indicates less-accessible regions. B) Proportions of genomic features annotated to chromatin peaks with differential accessibility after treatment with insulin or vehicle. C) Heatmap of differential gene expression in S2 cells after treatment with vehicle or insulin. Each row represents a significantly differentially-expressed transcript. The values in each row are scaled to the row mean. Red indicates an upregulated gene and blue indicates a downregulated gene.

Then, we investigated the correlation between all ATAC-seq and RNA-seq log_2_ fold changes after insulin treatment. The overall correlation between the two datasets was significant, but weak (*R* = 0.05, *p* = 1.8e-06; Fig. 4A). Performing the same analysis after stratifying by feature type showed correlations between the differential RNA-seq transcripts and the differential ATAC-seq features in addition to ≤1kb promoters (*R* = 0.096, *p* = 5.7e-13), including significant (though weak) correlations with ATAC-seq peaks in different promoter types (2-3 kb from the TSS: *R* = 0.15, *p* = 5.4e-4 and 1-2 kb from the TSS: *R* = 0.13, *p* = 9.0e-05). Furthermore, there were significant anticorrelations in ATAC-seq peaks for downstream elements (1-2 kb: *R* = -0.67, *p* = 0.035) and distal intergenic regions (*R* = -0.14, *p* = 0.0011; Fig. 4B). When we restricted the analysis to only the significant changes (FDR < 0.1) in ATAC-seq peaks, the overall correlation for all features increased 8-fold (*R* = 0.42, *p* = 4.2e-14; Fig. 5A), as did the correlations of several feature types (Fig. 5B). In particular, the correlation with promoters 1-2 kb from the TSS (*R* = 0.65, *p* = 1.3e-4) increased by approximately 5-fold. Conversely, the anticorrelations with peaks in downstream and distal intergenic regions disappeared. Together, these results suggest that DNA accessibility in distal promoters is involved in mediating changes in transcription.

**Figure 4.**
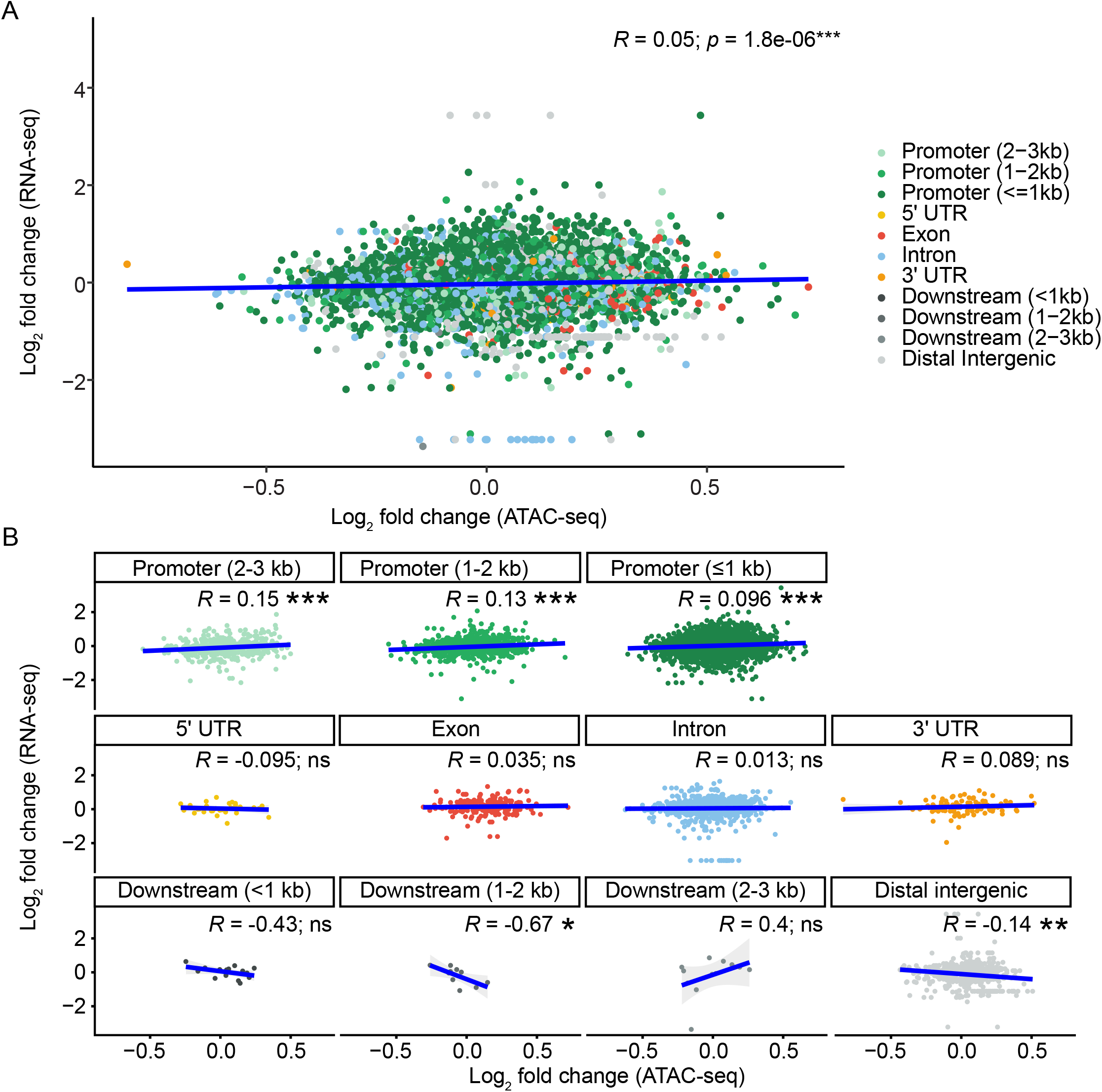
Insulin-induced log_2_ fold changes correlate between ATAC-seq and RNA-seq. A) Chromatin peaks were overlapped with expressed transcripts. Pearson correlation analysis shows a weak correlation between log_2_ fold changes in chromatin accessibility and transcript expression. B) Overlapping ATAC- and RNA-seq log_2_ fold change values from (A) were stratified by genomic feature. Pearson correlation analysis was used to identify correlations between ATAC- and RNA-seq counts by feature.

**Figure 5.**
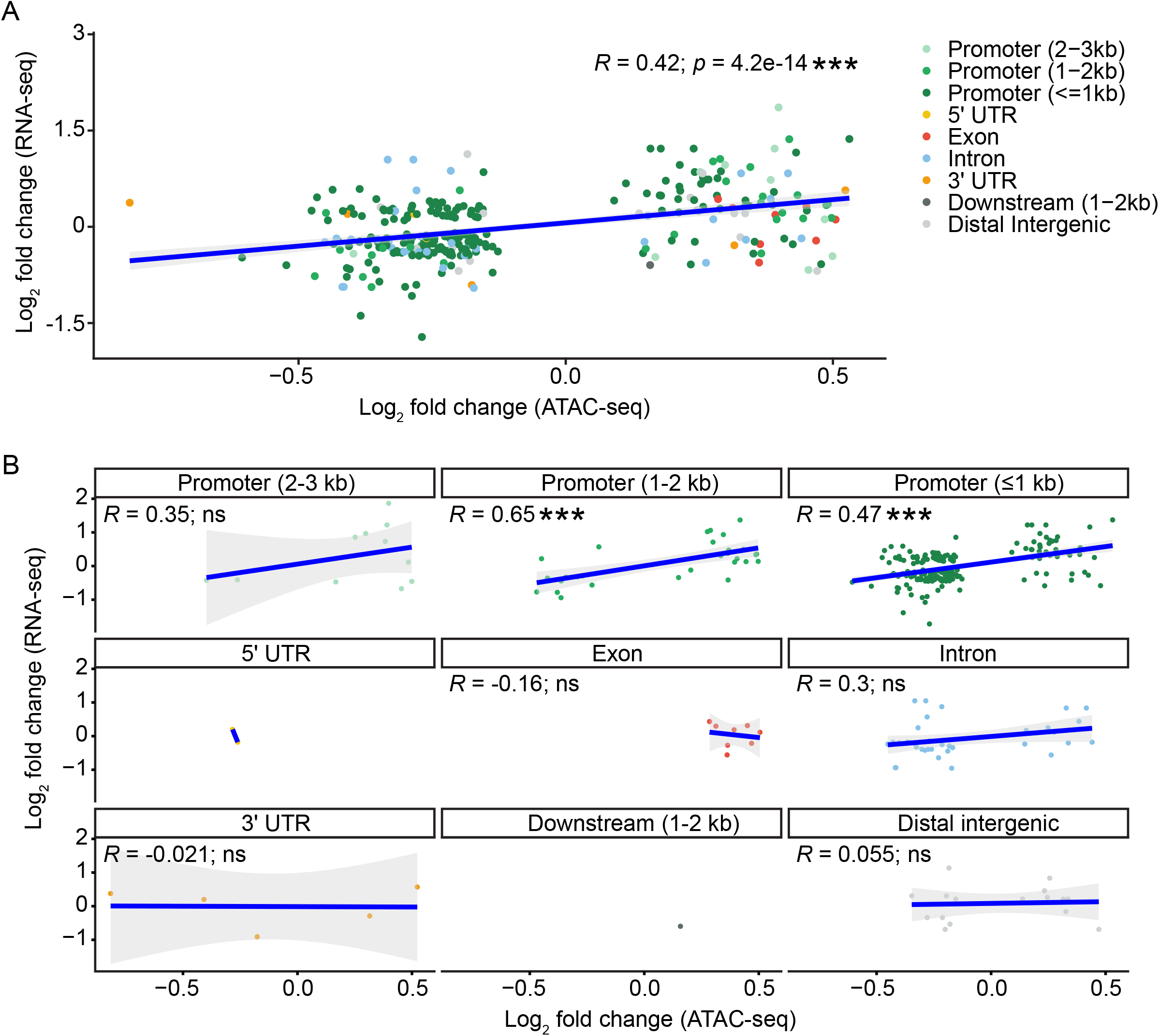
Significant insulin-induced changes in ATAC-seq indicates recruitment of distal promoters for transcript regulation. A) Significant Log_2_ fold change values from differentially-accessible chromatin peaks and differentially-expressed transcripts were analyzed by Pearson correlation analysis. B) Overlapping chromatin peaks and differentially-expressed genes from (A) were stratified by feature type and reveal distal (1-2 kb) promoter accessibility as correlated with insulin-induced transcript changes.

### Functional testing of significant differentially accessible ATAC-seq peaks

We next wanted to test whether any of the DNA regions from significantly more accessible ATAC-seq peaks could drive insulin-induced expression. We cloned a number of ATAC-seq peaks in front of a luciferase gene with a minimal promoter and transfected S2 cells with these vectors for 48 h. The cells were serum-starved for 18 h and then treated with 10 μM insulin or vehicle for 4 h. We selected three groups of four ATAC-seq peaks each: first, we chose the four peaks with the largest log_2_ fold change, indicating increased accessibility after insulin treatment (Additional File 1). Of the four tested plasmids, one showed significantly increased luciferase activity after insulin treatment: 3L114 (*p* = 0.016; Fig. 6A). Because ATAC-seq peaks in distal promoters were the most strongly correlated with differential gene expression in our above analysis (Fig. 5B), we next chose four peaks with the highest log_2_ fold change from distal promoter regions that were significantly more accessible after insulin. Of the tested peaks, 2 produced significantly increased luciferase activity after insulin treatment: 2L225 (*p* = 0.0033) and 2R111 (*p* = 0.025; Fig. 6B). Lastly, because introns often contain regulatory regions that contain instructive DNA for expression [18], we chose the four intron regions with the largest log_2_ fold changes for luciferase assays. One ATAC-seq peak resulted in significantly increased luciferase activity: X216 (*p* = 0.05) (Fig. 6C). These data show that DNA regions with increased accessibility upon insulin treatment can indeed drive insulin-induced increases in expression when placed in front of a heterologous promoter.

**Figure 6.**
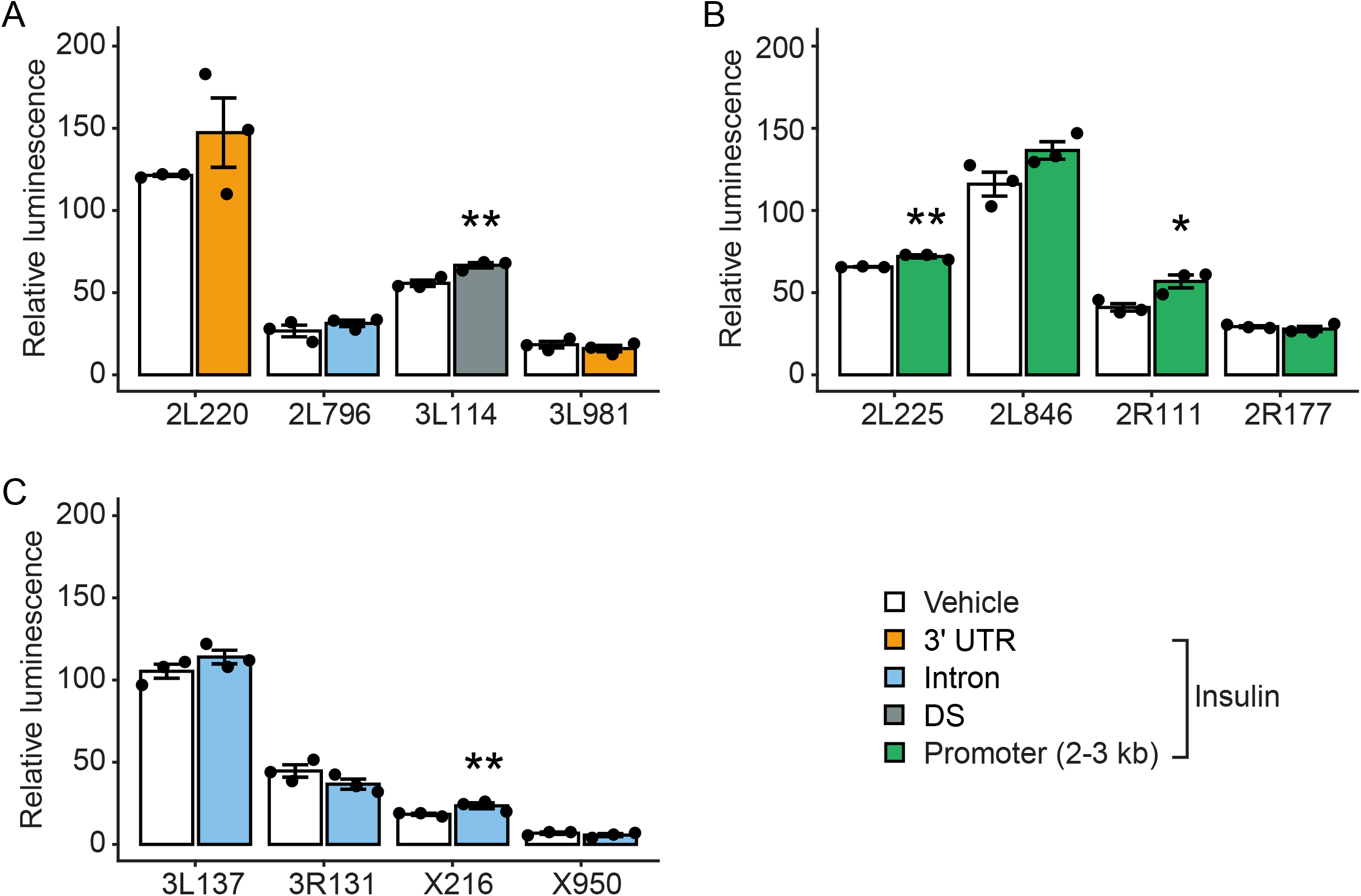
Cloned DNA from differentially-accessible chromatin regions can induce luciferase activity upon insulin application. DNA was cloned in front of a minimal promoter and luciferase gene, S2 cells were transfected for 18 h and then treated with insulin or vehicle for 4 h. A) Candidate ATAC-seq peaks were selected by the largest log_2_ fold change. B) Chromatin peaks from promoters (1-2 kb from the TSS) were the feature that was most highly correlated with differential transcript expression. Peaks with the highest log_2_ fold change from this correlation were cloned upstream of luciferase for functional validation. C) Chromatin peaks with significantly different accessibility were selected from introns, a genomic feature known to contain regulatory regions. Data represent means ± standard error of three biological replicates. Differences were analyzed by Student’s t-test.

### Limited predictability of the levels of expression and inducibility

Out of the twelve ATAC-seq peaks we cloned and tested, all led to significant – though variable – levels of luciferase expression, while only four caused significant insulin-inducibility. To determine whether the luciferase expression levels and inducibility by insulin was predictable from our ATAC-seq and RNA-seq datasets, we analyzed the correlation between the –omics data and luciferase activity. First, we asked if expression levels of luciferase were correlated with ATAC-seq peak reads, but found no correlation (Fig. 7A), even when we stratified the data according to distal promoter- (Fig. 7B) or intron-derived ATAC-seq peaks (Fig. 7C). Similarly, RNA-seq counts did not correlate with S2 luciferase luminescence (Fig. 7D-F). Next, we asked whether the log_2_ fold changes in the –omics data sets correlated with the relative inducibility of luciferase by insulin (measured as insulin/vehicle ratios). Again, we failed to observe significant correlations of S2 inducibility with log_2_ fold changes in ATAC-seq (Fig. 8A) and RNA-seq (Fig. 8B) reads, even when we analyzed only the cloned peaks that led to significant insulin-induced changes (Fig. 8C,D).

**Fig. 7.**
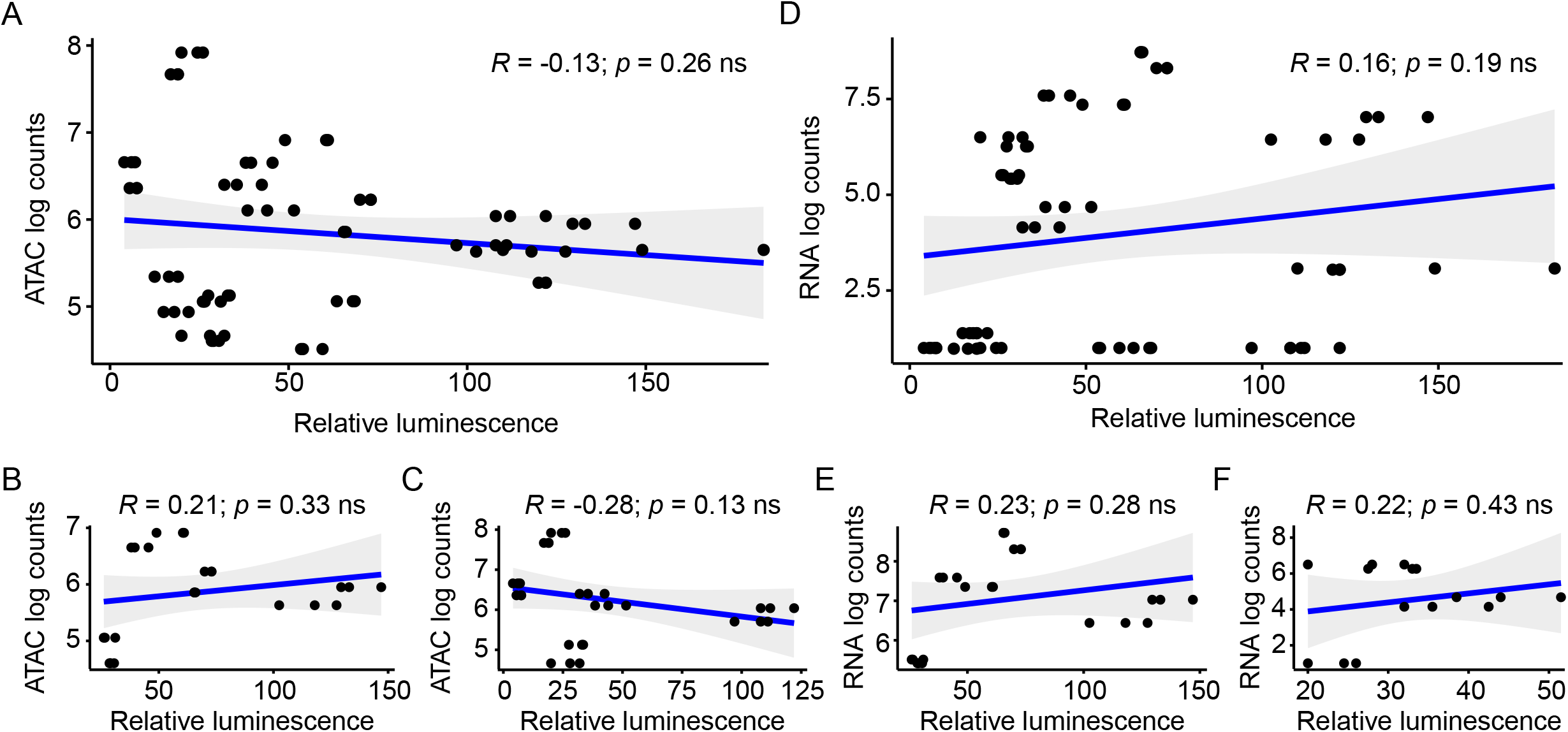
ATAC-seq and RNA-seq counts are not correlated with functional luciferase activity. Log-transformed ATAC-seq counts (A-C) and RNA-seq counts (D-F) from S2 cells were correlated to insulin-induced luciferase activity. A) Overall correlation between counts from the tested ATAC-seq peaks and luciferase activity; B) Promoters; C) Introns. D) Overall correlations between counts from genes annotated to the tested ATAC-seq peaks and luciferase activity; E) Promoters; F) Introns. Associations were analyzed by Pearson correlation analysis. Each point represents a biological replicate, and each peak was tested in triplicate.

**Fig. 8.**
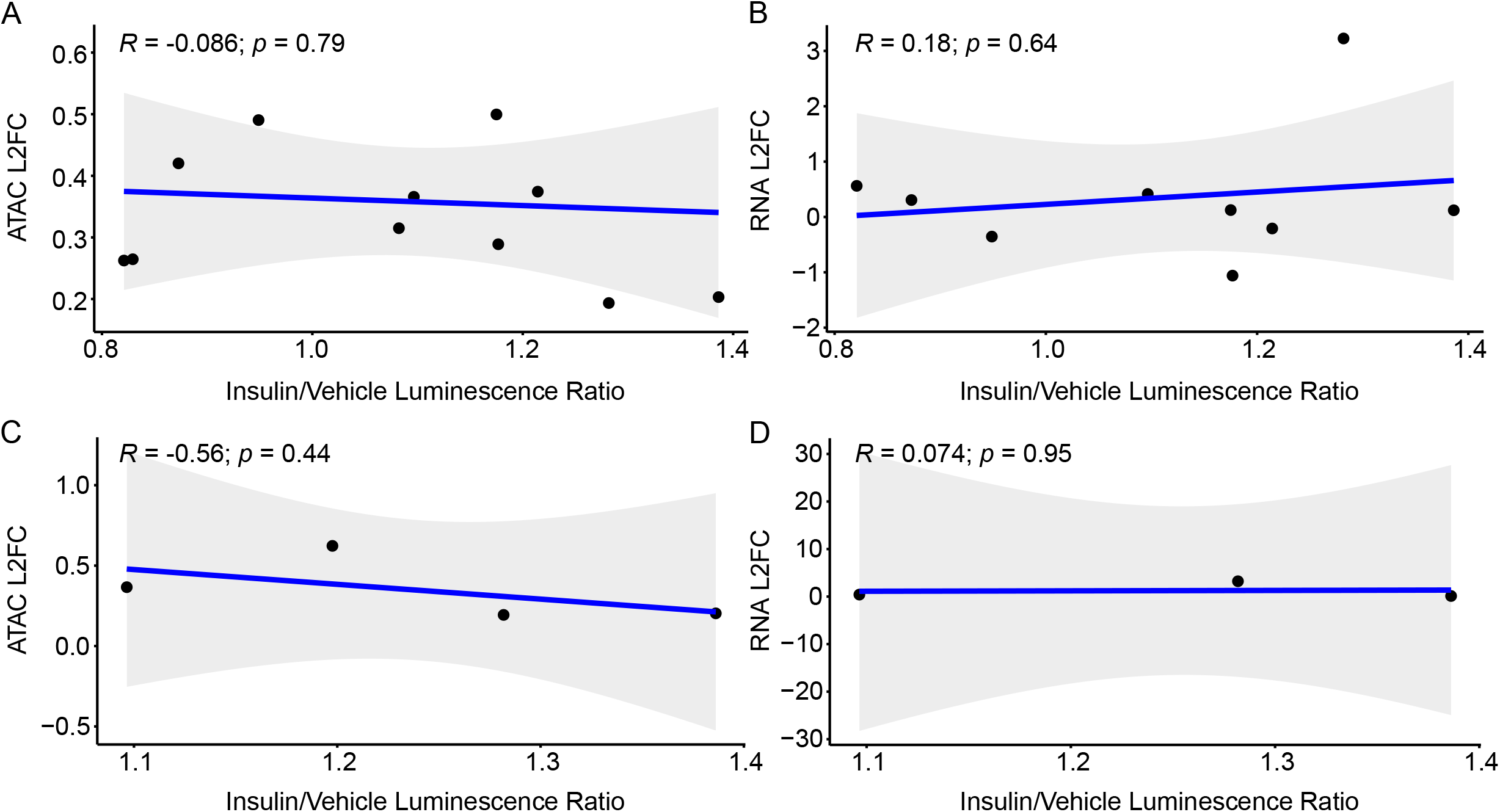
ATAC-seq and RNA-seq log_2_ fold changes do not predict insulin inducibility. A) ATAC-seq and B) RNA-seq log_2_ fold changes from S2 cells were correlated to insulin-induced luciferase activity, shown as the ratio between luciferase activity in vehicle- vs. insulin treated cells. C) Correlation between the ATAC-seq peaks driving significantly increased luciferase activity and associated ATAC-seq log_2_ fold changes. D) Correlation between the ATAC-seq peaks driving significantly increased luciferase activity and RNA-seq log_2_ fold changes in the associated genes. Associations were analyzed by Pearson correlation analysis. Each point represents a biological replicate, and each peak was tested in triplicate.

## DISCUSSION

Next-generation sequencing has enabled an unprecedented amount of genome-wide information on RNA transcript levels and DNA accessibility. ATAC-seq data provides accessibility information from distinct features/regions of a gene, thereby suggesting gene regions that act as functional enhancers (or suppressors) of gene expression. Here, we investigated the correlation between genome-wide changes in DNA accessibility and transcript levels and found significant correlations that were mostly driven by proximal and distal promoter regions. Cloning some of these DNA regions with increased accessibility upon insulin stimulation showed that some of them indeed act as transcriptional enhancers, demonstrating that genome-wide ATAC-seq can be harnessed to clone functionally-active insulin-response elements.

To investigate the functional relevance of differential DNA accessibility, we first determined genome-wide ATAC-seq reads in *Drosophila* S2 cells from serum-starved and insulin-exposed conditions (Fig. 1). The insulin receptor activates several downstream pathways, including the PI3K [19] and Ras/ERK [20] pathways, which have various effects on the chromatin state [21, 22] and gene expression [23] during several cellular processes including cell growth, protein synthesis, and gluconeogenesis [24]. Thus, activating insulin signaling provided a way to identify broad chromatin and gene expression changes, which allowed us to integrate these physiological changes and determine whether chromatin regions that become more open after insulin signaling could predict gene regulation. We found significant overall correlations between ATAC-seq reads and transcript levels, driven by ATAC-seq peaks in proximal promoters (Fig. 2). In ATAC-seq, genome regions with increased accessibility result in a higher mapped read count [9]. Because promoter regions are critical for the initiation of transcription, these genomic regions are generally accessible for actively-transcribed genes [25]. Thus, proximal promoter regions largely drive the overall correlation between ATAC-seq and RNA-seq counts that we observed. These data indicate that normalized counts can identify correlations between chromatin and gene expression, but these correlations are likely limited to promoter regions for actively transcribed genes.

When we analyzed correlations between all insulin-induced log_2_ fold changes in ATAC-seq peak and transcript reads (Fig. 3), changes in open chromatin in distal (1-2 and 2-3 kb away from the TSS) promoter regions also correlated significantly with changes in transcript levels (Fig. 4). This suggests that the application of insulin recruits additional distal promoters that participate in promoting transcription. Conversely, other distal promoter regions become less accessible, and the linked genes are less transcribed with insulin. These distal and proximal promoter correlations with transcript levels became even stronger when we only analyzed ATAC-seq peaks that changed significantly with insulin (Fig. 5). These results suggest that perturbations that cause differential gene expression occur via recruitment of additional regulatory promoter features. The correlations between differential transcript levels and differentially accessible promoter regions were all positive, suggesting that these regions play a role in the downstream differential gene expression. However, these data do not exclude the possibility that in some genes, insulin might lead to increased accessibility at promoters which are then bound by transcriptional repressors, leading to decreased transcription. Indeed, numerous ATAC-seq peak/transcript data points are in quadrants of anticorrelation (Fig. 5), and the insulin-induced transcription factor FOXO is known to have transcriptional repressor activity [26, 27]. Future experiments focusing on such anticorrelated data pairs/genes might reveal DNA regions that lead to insulin-induced transcriptional repression.

Our main goal was to determine whether we could harness our ATAC-seq data to generate insulin-inducible reporter plasmids. We selected ATAC-seq peaks based on our correlation analysis of differentially-accessible chromatin regions and differential transcript expression. We particularly focused on more distal promoters (1-2 kb from the TSS) because the correlation increase was the largest for this feature. Distal promoter regions may include regulatory regions such as enhancers or repressors that are critically involved in regulating gene expression [28]. Our results suggested that these regions can drive differential gene expression (Fig. 6). We also selected peaks with relatively large log_2_ fold changes in intron peaks. In *Drosophila*, intronic regions often contain regulatory sequences [18], thus altering chromatin accessibility in genome regions associated with introns is one mechanism to control gene expression [29]. Finally, we selected peaks with the largest log_2_ fold changes, irrespective of feature type. In each of these three categories we found peaks that led to significant insulin-induced increases in reporter gene expression. However, none of the three categories seemed obviously more promising for predicting insulin-inducibility. Furthermore, neither read counts nor log_2_ fold changes in ATAC-seq or RNA-seq were predictive of insulin-inducibility (Figs. 7, 8). This suggests that while ATAC-seq data can be successfully harnessed to generate insulin-inducible reporter constructs, their efficacy is not obviously predictable and will require larger datasets to understand which ATAC-seq peaks can be utilized to generate functionally relevant transgenes. Indeed, previous studies investigating putative enhancer elements identified candidates based on overlap with known histone marks (H3K4me1, H3K27ac, etc. identified by ChIP-seq), known enhancers associated with annotated genes of interest [30–32], or used massively parallel reporter assays [33]. In contrast, our goal was to determine whether ATAC-seq alone could predict downstream transcription using on feature-based or fold change-based selection. Importantly, these previous studies showed similar success rates to ours. Peaks with increased chromatin accessibility after insulin treatment that did not result in insulin-induced luciferase activity may represent regulatory elements that are involved in setting up poised transcription or may contain repressor regions that pause transcription. In contrast, peaks causing increased luciferase activity may represent sequences that are sufficient to initiate transcription or activate promoter clearance [34–36]. Additional studies using ChIP-seq to identify the histone marks at our tested peak sequences will be required to determine whether they are enhancers involved in poised versus active transcription.

## CONCLUSIONS

Our investigation shows that ATAC-seq data can be harnessed to isolate regulatory DNA regions that are both expressed and inducible. However, because chromatin peaks may be one of several regulatory sequences [18, 28], these chromatin regions cannot be easily predicted by analysis of these genome-wide –omics data alone and must be functionally validated. Still, our data show the feasibility of using ATAC-seq to generate active transgenes that are inducible by an intervention or by a diseased state to drive a reporter, or even a disease-antidote gene.

## METHODS

### Cell culture

Drosophila S2 cells (Drosophila Genomics Resource Center, Bloomington, IN, USA) were cultured in Schneider’s Drosophila Medium (ThermoFisher, Waltham, MA, USA) supplemented with 10% fetal bovine serum (ThermoFisher) at 25 °C. Cells were cultured in Schneider’s medium without FBS for 24 h before experiments. Then, cells were incubated with 10 μM insulin (Sigma Aldrich, St. Louis, MO, USA) or vehicle (25 mM HEPES, pH 8.2) for 4 h at 25 °C.

### ATAC-seq

S2 cells were incubated with 3 μM DAPI for 10 min. 60,000 cells per sample were sorted using a BD FACS Aria flow cytometer (BD Biosciences, San Jose, CA, USA). DAPI-negative cells were collected into ice-cold PBS (pH 7.4). After sorting, the samples were washed once with ice-cold PBS and centrifuged at 500 g for 5 min at 4 °C. ATAC-seq libraries were prepared as previously described [37]. Briefly, 50 μL lysis buffer (10 mM Tris-HCl 7.4, 10 mM NaCl, 3 mM MgCl2, 0.1% NP40) was added to each sample, and the sample was centrifuged at 500 g for 10 min at 4 °C. The supernatant was removed, and the nuclei pellet was tagmented using a Nextera DNA Library Prep kit (Illumina, Inc., San Diego, CA, USA) as previously described. Then, the tagmented DNA was purified using a Qiagen MinElute PCR Purification Kit (Qiagen, Germantown, MD, USA). The purified DNA was PCR amplified for 5 cycles using a Nextera DNA Library Index kit (Illumina) and Phusion HF Master Mix (New England BioLabs, Inc., Ipswich, MA, USA) with the following protocol: 72 °C for 5 min, 98 °C for 30 sec, and 5 cycles of 98 °C for 10 sec, 63 °C for 30 sec, and 72 °C for 1 min. A 5-μL aliquot of the pre-amplified reaction was analyzed by qPCR using SsoFast EvaGreen Supermix (Bio-Rad Life Science, Inc., Hercules, CA, USA) and an Applied Biosystems 7900HT qPCR instrument (ThermoFisher) using the following protocol: 1 cycle of 98 °C for 30 sec and 40 cycles of 98 °C for 10 sec, 63 °C for 30 sec, and 72 °C for 1 min. Then, the pre-amplified PCR mixture was amplified for another 10 cycles (corresponding to 1/3 maximum fluorescence from the qPCR assay) using the same thermocycling parameters. After amplification, the libraries were purified using AMPure XP beads (Beckman Coulter Life Sciences, Indianapolis, IN, USA). Libraries were sequenced on an Illumina HiSeq 2500 instrument using 50-bp single-end reads.

### RNA-seq

Total RNA was isolated from S2 cells using a PureLink RNA purification kit (ThermoFisher). Then, rRNA was removed from each sample with a Ribo-Zero rRNA Removal kit (Illumina). RNA libraries were constructed using a NEBNext Ultra II RNA Library Kit for Illumina and NEBNext Multiplex Oligos for Illumina, Primer Set 1 (New England Biolabs). Libraries were sequenced on an Illumina HiSeq 2500 instrument using 50-bp single-end reads.

### ATAC-seq data analysis

ATAC-seq Fastq files were aligned to the dm6 genome assembly (http://ftp.flybase.net/releases/FB2018_06/dmel_r6.25/fasta/) using Novocraft Novoalign with the following settings: --NonC -o SAM -r Random. SAM files were processed to BAM format, sorted, and indexed using Samtools [38]. BAM files were reads per million-normalized and converted to bigWig files using the Bio-ToolBox ‘bam2wig.pl’ program (https://github.com/tjparnell/biotoolbox/blob/master/scripts/bam2wig.pl). Peak calling was performed on the bigWig files by utilizing the Multi-Replicate Macs ChIPseq pipeline (https://github.com/HuntsmanCancerInstitute/MultiRepMacsChIPseq) with the following settings: --dupfrac 0.2 --size 200 --cutoff 2 --peaksize 300 --peakgap 200. Called peaks were annotated in R with the ChIPseeker package [39]. Count data for called peaks was collected from processed BAM files using the Bio-ToolBox ‘get_datasets.pl’ program (https://metacpan.org/pod/distribution/Bio-ToolBox/scripts/get_datasets.pl). The count data was then used to identify differentially accessible regions with the R package DEseq2 [40].

### RNA-seq data analysis

RNA-seq fastq files were aligned to the BDGP6 genome assembly using the STAR aligner [41] with the following settings: --twopassMode Basic --outSAMtype BAM SortedByCoordinate -- outWigType bedGraph --outWigStrand Unstranded --clip3pAdapterseq AGATCGGAAGAGCACACGTCTGAACTCCAGTCA. The resulting sorted BAM files were indexed using Samtools (Li et al., 2009). FeatureCounts was used to collect count data for BDGP6 genes using the following command: -T 16 -s 2 [42]. Count data for all replicates and experimental conditions were combined into a single count matrix in R. The count matrix was subsequently used to identify differentially expressed genes with the R package DEseq2 [40].

### Integration analysis of ATAC-Seq and RNA-Seq datasets

The ATAC-seq peak data were compared to the RNA-seq data to determine how chromatin accessibility influenced gene expression. The raw RNA-seq and ATAC-seq counts for each sample were compared using the gene annotation of the ATAC-seq peak and the assigned RNA-seq gene. The raw count value was averaged by experimental condition and genomic assay type. Then, the RNA-seq and ATAC-seq datasets were compared using the annotated genes and the log_2_ fold change values for each peak/gene in the respective genomic assay. ATAC-seq peaks with an FDR < 0.1 and genes detected by RNA-seq with an FDR < 0.05 were used to compare the differentially accessible peaks and differentially expressed genes. Pearson correlation analysis was performed between the log_2_ fold change values of the genomic assays and between the average raw count values of the genomic assays (controlling for the experimental condition).

### Plasmid construction and transformation

A multiple cloning site (MCS) was cloned into the backbone pDEST VanGlow-GL vector [43]. Then, we removed the *mini-white*^*+*^ cassette using the restriction enzymes AflII (3L137, 3R131, X216, 3L114, 2L796, 3L981, 2L220, 2R177, 2L225, and FOXO TFBS) or SmaI and PmlI (X950, 2L846, and 2R111). The digested plasmids were incubated with T4 ligase for 15 min at room temperature and purified by 1% gel electrophoresis. The plasmids were linearized using AvrII and PacI (sites contained in MCS; all enzymes from New England BioLabs).

Genomic DNA was purified from S2 cells using a Monarch Genomic DNA Purification kit. Candidate chromatin peak sequences and 100-200 bp flanking sequences [32] (Additional File 1) were amplified using Phusion High-Fidelity PCR MasterMix (primer sequences are listed in Additional File 2) and a C1000 thermocycler (Bio-Rad Life Science). The peak sequences were amplified for 98 °C for 5 min, followed by 35 cycles of 98 °C for 30 sec, 52-68 °C gradient for 30 sec, 72 °C for 4 min, and a final incubation for 5 min at 72 °C. The amplified fragments were purified on 1% agarose gels and extracted using a Monarch Gel Purification kit and cloned into linearized VanGlo-GL-MCS plasmid using NEBuilder HiFi Assembly master mix. The assembled plasmids were transformed into DH5α cells and grown overnight. Clones were screened by restriction digestion using EcoRI-HF. Sequences were confirmed by Sanger sequencing at GeneWiz (South Plainfield, NJ, USA). Confirmed plasmids were transformed into S2 cells using TransIT Insect Transfection Reagent (Mirus Bio, Madison, WI, USA). Transformed cells were grown for 48 h before use in experiments.

### Luciferase assays

Transformed S2 cells were serum starved for 24 h and treated with insulin or vehicle as described above (*Cell culture*). Then, luciferase activity was assayed using a Luciferase Reporter Substrate Assay Kit-Firefly (Abcam, Cambridge, MA, USA). Luminescence was detected with a BioTek Synergy HTX microplate reader (BioTek Instruments, Winooski, VT, USA) and Gen5 2.01.17 software (BioTek Instruments).

### Statistical analysis

Statistical differences in relative luminescence data were analyzed by Student’s t-tests with at least three biological replicates using GraphPad Prism 9.0 software. Differences between genome feature proportions were analyzed using χ^2^ tests included in R [44] Correlations were analyzed using Pearson correlation tests included in R. Heatmaps were generated using the R package ComplexHeatmap [45]. PCA plots were created using the R package pcaExplorer [46]. Correlation plots were produced with the R package ggpubr (https://github.com/kassambara/ggpubr).

## DECLARATIONS

### Ethics approval and consent to participate

Not applicable

### Consent for publication

Not applicable

### Availability of data and materials

All sequencing data are deposited in the Sequence Read Archive (BioProject ID: PRJNA730574; https://www.ncbi.nlm.nih.gov/sra/PRJNA730574). Plasmids developed in this study are available upon reasonable request.

### Author contributions

CBM performed the ATAC-seq and RNA-seq experiments, constructed the luciferase plasmids, analyzed data, and wrote the manuscript. ABM analyzed the sequencing data and wrote the manuscript. MAP constructed luciferase plasmids and revised the manuscript. ARR edited the manuscript and procured funding. AR oversaw the project, edited the manuscript, and procured funding. All authors read and approved the final manuscript.

## Acknowledgments

We thank members of the lab for continued discussion. This work was supported by the University of Utah Flow Cytometry Facility, the University of Utah Genomics Core Facility, the High Throughput Sequencing Core at the Huntsman Cancer Institute, and the National Cancer Institute through Award Number 5P30CA042014. The content is solely the responsibility of the authors and does not necessarily represent the official views of the National Institutes of Health.

## Additional files

Additional_File_1.xslx; ATAC-seq peaks that were cloned and tested in luciferase assays.

Additional_File_2.xslx; Primer sequences used for cloning.

## REFERENCES

1. MacNeil LT, Walhout AJM. Gene regulatory networks and the role of robustness and stochasticity in the control of gene expression. Genome Research. 2011;21:645–57.

2. Goode DK, Obier N, Vijayabaskar MS, Lie-A-Ling M, Lilly AJ, Hannah R, et al. Dynamic Gene Regulatory Networks Drive Hematopoietic Specification and Differentiation. Dev Cell. 2016;36:572–87.

3. Tokusumi Y, Tokusumi T, Shoue DA, Schulz RA. Gene regulatory networks controlling hematopoietic progenitor niche cell production and differentiation in the Drosophila lymph gland. PLoS One. 2012;7:41604.

4. Morozova T V, Mackay TFC, Anholt RRH. Transcriptional networks for alcohol sensitivity in Drosophila melanogaster. Genetics. 2011;187:1193–205.

5. Duarte FM, Fuda NJ, Mahat DB, Core LJ, Guertin MJ, Lis JT. Transcription factors GAF and HSF act at distinct regulatory steps to modulate stress-induced gene activation. 2016.

6. Petruccelli E, Brown T, Waterman A, Ledru N, Kaun KR. Alcohol Causes Lasting Differential Transcription in Drosophila Mushroom Body Neurons. 2020.

7. Jothi R, Cuddapah S, Barski A, Cui K, Zhao K. Genome-wide identification of in vivo protein-DNA binding sites from ChIP-Seq data. Nucleic Acids Res. 2008;36:5221–31.

8. Song L, Crawford GE. DNase-seq: A high-resolution technique for mapping active gene regulatory elements across the genome from mammalian cells. Cold Spring Harb Protoc. 2010;5:pdb.prot5384.

9. Buenrostro JD, Giresi PG, Zaba LC, Chang HY, Greenleaf WJ. Transposition of native chromatin for fast and sensitive epigenomic profiling of open chromatin, DNA-binding proteins and nucleosome position. Nat Methods. 2013;10:1213–8.

10. Kagohara LT, Zamuner F, Davis-Marcisak EF, Sharma G, Considine M, Allen J, et al. Integrated single-cell and bulk gene expression and ATAC-seq reveals heterogeneity and early changes in pathways associated with resistance to cetuximab in HNSCC-sensitive cell lines. Br J Cancer 2020 1231. 2020;123:101–13.

11. Li X, Chen Y, Fu C, Li H, Yang K, Bi J, et al. Characterization of epigenetic and transcriptional landscape in infantile hemangiomas with ATAC-seq and RNA-seq. https://doi.org/102217/epi-2020-0060. 2020;12:893–905.

12. Ackermann AM, Wang Z, Schug J, Naji A, Kaestner KH. Integration of ATAC-seq and RNA-seq identifies human alpha cell and beta cell signature genes. Mol Metab. 2016;5:233– 44.

13. McNeill H, Craig GM, Bateman JM. Regulation of neurogenesis and epidermal growth factor receptor signaling by the insulin receptor/target of rapamycin pathway in drosophila. Genetics. 2008;179:843–53.

14. Kido Y, Nakae J, Accili D. The Insulin Receptor and Its Cellular Targets 1. J Clin Endocrinol Metab. 2001;86:972–9.

15. Puig O, Marr MT, Ruhf ML, Tjian R. Control of cell number by Drosophila FOXO: downstream and feedback regulation of the insulin receptor pathway. 2003.

16. Kulkarni MM, Kulkarni MM, Sopko R, Sun X, Hu Y, Nand A, et al. An Integrative Analysis of the InR/PI3K/Akt Network Identifies the Dynamic Response to Insulin Signaling. Cell Rep. 2016;16:3062–74.

17. Post S, Karashchuk G, Wade JD, Sajid W, De Meyts P, Tatar M. Drosophila Insulin-Like Peptides DILP2 and DILP5 Differentially Stimulate Cell Signaling and Glycogen Phosphorylase to Regulate Longevity. Front Endocrinol (Lausanne). 2018;9:245.

18. Roy S, Ernst J, Kharchenko P V., Kheradpour P, Negre N, Eaton ML, et al. Identification of functional elements and regulatory circuits by Drosophila modENCODE. Science (80-). 2010;330:1787–97.

19. Dekanty A, Lavista-Llanos S, Irisarri M, Oldham S, Wappner P. The insulin-PI3K/TOR pathway induces a HIF-dependent transcriptional response in Drosophila by promoting nuclear localization of HIF-α /Sima. J Cell Sci. 2005;118:5431–41.

20. Zhang W, Thompson BJ, Hietakangas V, Cohen SM. MAPK/ERK signaling regulates insulin sensitivity to control glucose metabolism in Drosophila. PLoS Genet. 2011;7:1002429.

21. Mouchel-Vielh E, Rougeot J, Decoville M, Peronnet F. The MAP kinase ERK and its scaffold protein MP1 interact with the chromatin regulator Corto during Drosophila wing tissue development. BMC Dev Biol. 2011;11:1–14.

22. Sánchez-Alegría K, Flores-León M, Avila-Muñoz E, Rodríguez-Corona N, Arias C. PI3K signaling in neurons: A central node for the control of multiple functions. International Journal of Molecular Sciences. 2018;19:3725.

23. Garofalo RS. Genetic analysis of insulin signaling in Drosophila. Trends in Endocrinology and Metabolism. 2002;13:156–62.

24. Goberdhan DCI, Wilson C. The functions of insulin signaling: Size isn’t everything, even in Drosophila. Differentiation. 2003;71:375–97.

25. Blythe SA, Wieschaus EF. Establishment and maintenance of heritable chromatin structure during early Drosophila embryogenesis. Elife. 2016;5.

26. Jünger MA, Rintelen F, Stocker H, Wasserman JD, Végh M, Radimerski T, et al. The Drosophila Forkhead transcription factor FOXO mediates the reduction in cell number associated with reduced insulin signaling. J Biol. 2003;2:20.

27. Ramaswamy S, Nakamura N, Sansal I, Bergeron L, Sellers WR. A novel mechanism of gene regulation and tumor suppression by the transcription factor FKHR. Cancer Cell. 2002;2:81–91.

28. Gisselbrecht SS, Palagi A, Kurland J V., Rogers JM, Ozadam H, Zhan Y, et al. Transcriptional Silencers in Drosophila Serve a Dual Role as Transcriptional Enhancers in Alternate Cellular Contexts. Mol Cell. 2020;77:324-337.e8.

29. Duret L. Why do genes have introns? Recombination might add a new piece to the puzzle. Trends in Genetics. 2001;17:172–5.

30. Daugherty AC, Yeo RW, Buenrostro JD, Greenleaf WJ, Kundaje A, Brunet A. Chromatin accessibility dynamics reveal novel functional enhancers in C. elegans. Genome Res. 2017;27:2096–107.

31. Quillien A, Abdalla M, Yu J, Ou J, Zhu LJ, Lawson ND. Robust Identification of Developmentally Active Endothelial Enhancers in Zebrafish Using FANS-Assisted ATAC-Seq. Cell Rep. 2017;20:709–20.

32. Cusanovich DA, Reddington JP, Garfield DA, Daza RM, Aghamirzaie D, Marco-Ferreres R, et al. The cis-regulatory dynamics of embryonic development at single-cell resolution. Nature. 2018;555:538–42.

33. Hrvatin S, Tzeng CP, Nagy MA, Stroud H, Koutsioumpa C, Wilcox OF, et al. A scalable platform for the development of cell-type-specific viral drivers. Elife. 2019;8.

34. Klemm SL, Shipony Z, Greenleaf WJ. Chromatin accessibility and the regulatory epigenome. Nature Reviews Genetics. 2019;20.

35. Bae S, Lesch BJ. H3K4me1 Distribution Predicts Transcription State and Poising at Promoters. Front Cell Dev Biol. 2020;8:289.

36. Koenecke N, Johnston J, Gaertner B, Natarajan M, Zeitlinger J. Genome-wide identification of Drosophila dorso-ventral enhancers by differential histone acetylation analysis. Genome Biol. 2016;17:1–19.

37. Buenrostro JD, Wu B, Chang HY, Greenleaf WJ. ATAC-seq: A method for assaying chromatin accessibility genome-wide. Curr Prot Mol Biol. 2015 Jan;109(1):21–9.

38. Li H, Handsaker B, Wysoker A, Fennell T, Ruan J, Homer N, et al. The Sequence Alignment/Map format and SAMtools. Bioinformatics. 2009;25:2078–9.

39. Yu G, Wang LG, He QY. ChIP seeker: An R/Bioconductor package for ChIP peak annotation, comparison and visualization. Bioinformatics. 2015;31:2382–3.

40. Anders S, Huber W. Differential expression analysis for sequence count data. 2010.

41. Dobin A, Davis CA, Schlesinger F, Drenkow J, Zaleski C, Jha S, et al. STAR: Ultrafast universal RNA-seq aligner. Bioinformatics. 2013;29:15–21.

42. Liao Y, Smyth GK, Shi W. FeatureCounts: An efficient general purpose program for assigning sequence reads to genomic features. Bioinformatics. 2014;30:923–30.

43. Janssens DH, Hamm DC, Anhezini L, Xiao Q, Siller KH, Siegrist SE, et al. An Hdac1/Rpd3-Poised Circuit Balances Continual Self-Renewal and Rapid Restriction of Developmental Potential during Asymmetric Stem Cell Division. Dev Cell. 2017;40:367-380.e7.

44. R Core. R.:A language and environment for statistical computing. 2020.

45. Gu Z, Eils R, Schlesner M. Complex heatmaps reveal patterns and correlations in multidimensional genomic data. Bioinformatics. 2016;32:2847–9.

46. Marini F, Binder H. PcaExplorer: An R/Bioconductor package for interacting with RNA-seq principal components. BMC Bioinformatics. 2019;20:1–8.

